# A F_420_-dependent single domain chemogenetic tool for protein de-dimerization

**DOI:** 10.1101/2022.11.07.515376

**Authors:** James Antoney, Stephanie Kainrath, F. Hafna Ahmed, Suk Woo Kang, Emily R. R. Mackie, Tatiana P. Soares da Costa, Colin J. Jackson, Harald Janovjak

## Abstract

Protein-protein interactions (PPIs) mediate many fundamental cellular processes and their control through optically or chemically responsive protein domains has a profound impact on basic research and some clinical applications. Most available chemogenetic methods induce the association, i.e., dimerization or oligomerization, of target proteins, and the few available dissociation approaches either break large oligomeric protein clusters or heteromeric complexes. Here, we have exploited the controlled dissociation of a dimeric oxidoreductase from mycobacteria (MSMEG_2027) by its native cofactor, F_420_, which is not present in mammals, as a bioorthogonal monomerization switch. We found that in the absence of F_420_, MSMEG_2027 forms a unique domain-swapped dimer that occludes the cofactor binding site. Substantial remodelling of the intertwined N-terminal helix upon F_420_ binding results in the dissolution of the dimer. We then show that MSMEG_2027 can be expressed as fusion proteins in human cells and apply it as a tool to induce and release MAPK/ERK signalling downstream of a chimeric fibroblast growth factor receptor 1 (FGFR1) tyrosine kinase. This F_420_-dependent chemogenetic de-dimerization tool is stoichiometric, based on a single domain and presents a novel mechanism to investigate protein complexes *in situ*.

## INTRODUCTION

The dynamic formation and disruption of PPIs is a core mechanism that underlies the regulation of a wide range of cellular processes, including subcellular protein localisation, enzymatic activity, recognition of extracellular signals, downstream signal transduction and gene transcription. To establish causal linkages between PPIs and cell and organism behaviour, genetically encoded synthetic biology tools have been developed that facilitate both the observation as well as the regulation of PPIs. In particular, chemogenetic tools utilise small molecules as interaction regulators, which can be applied over prolonged periods of time, whereas optogenetic tools respond to light triggers, which are beneficial for fast and local control. A number of available chemogenetic tools to induce protein homo- or heterodimerization have now been applied in dozens of *in vitro* and *in vivo* studies, including on signalling dynamics [1-4], protein expression [5], enzyme activity [6, 7], histone modification [8], and even as chemically inducible “kill switches” in human cell therapy [9, 10].

While methods to *induce* PPIs of target proteins are established, few effective tools are available for *disrupting* PPIs with high fidelity. The F36M variant of the FK506 binding protein (FKBP) has been employed for ligand-reversible aggregation [11]. At least three sequential domain copies are required for significant interaction in cells, and such fusion proteins aggregate into oligomers of variable higher-order stoichiometry [11, 12]. Another example is the reversible chemical dimerizer rCD1, which is a bimodal synthetic ligand of FKBP. Here, a protein of interest tagged with FKBP is, upon addition of rCD1, recruited and covalently bound to a SNAP-tagged protein partner [13]. FK506 can then be added to compete with rCD1 for binding to FKBP resulting in the release of the interaction [14]. The use of both FKBP and a SNAP tag for the rCD1 system prevents their use in tandem experimental designs and the system consists of four components (two protein domains and two ligands) with differential (non)covalent binding kinetics. In conceptually related approaches, monovalent drugs displace divalent crosslinkers of the N-terminal domains of bacterial Gyrase B [15-17] or FKBP-based systems [18]. More recently, elegant work demonstrated disruption of the interactions between the viral NS3a protease or members of the Bcl-2 protein family and cognate peptides using clinical PPI inhibitors [19-21]. The peptides are either fused to target proteins [19] or grafted onto designer protein domains [20, 21]. Remarkably, displacement of monovalent binders has been very recently demonstrated to result in the reversible oligomerization the BTB domain of the transcription factor BCL6 [22]. Whilst there has been exciting recent progress, existing approaches to break PPIs utilize synthetic small molecules, rely on competitive displacement and/or are limited to heteromeric interactions or large complexes; consequently, recent systems also require at least three molecular components (one/two protein domain(s) and/or small molecule(s)).

Here, we present an inspired-by-nature tool for chemogenetic de-dimerization (CDD) based on the flavin/deazaflavin oxidoreductase (FDOR) MSMEG_2027 from *Mycobacterium smegmatis*. MSMEG_2027 monomerizes upon binding of its native enzyme cofactor F_420_, a deazaflavin that is exclusively synthesized by archaea and a subset of bacteria [23-25]. We structurally characterized the dimeric apoenzyme and monomeric cofactor-bound state, as well as the thermodynamics of F_420_ binding. We then employed MSMEG_2027 to induce dimerization and constitutive signalling activity in a prototypical receptor tyrosine kinase and show modulation of cell signalling by F_420_ treatment in human cells, demonstrating the potential of F_420_:MSMEG_2027 as a high affinity, single domain bioorthogonal CDD tool.

## RESULTS

### F_420_ binding monomerizes MSMEG_2027

MSMEG_2027 belongs to the FDOR-A subgroup of the FDOR superfamily and has been shown to function as an F_420_H_2_-dependent quinone reductase [23, 26]. Because the native activity requires reduction of F_420_ by F_420_-dependent glucose-6-phosphate dehydrogenase (FGD1; not present in eukaryotes [27]), this enzyme will not be functional as a quinone reductase in recombinant eukaryotic systems. F_420_ from *M. smegmatis* (predominantly F_420_-5,6,7, indicating the presence of five to seven glutamate residues in the side chain) binds MSMEG_2027 with high affinity (*K*_D_ = 0.5 ± 0.1 μM) and high specificity; the structurally similar and ubiquitous flavin cofactor flavin mononucleotide (FMN) has ∼50-fold lower binding affinity [23]. The structure of a truncated form of the apo-MSMEG_2027 lacking the first 26 amino acids (NΔ26-MSMEG_2027) was previously solved to a resolution of 1.5 Å (PDB ID 4Y9I) and confirmed that the protein adopts a split β-barrel fold, albeit the truncated form was found to be inactive [23].

During purification of the full-length construct of MSMEG_2027, we observed by size exclusion chromatography that, unlike the truncated protein, full-length apo-MSMEG_2027 is dimeric in solution **(Supplementary Figure S1a**,**b)**. We then found that complete dissociation of the dimer could be induced by addition of an excess of F_420_ **(Supplementary Figure S1c)**. Addition of limiting amounts of F_420_ resulted in two peaks corresponding to the monomeric and dimeric forms, of which only the peak corresponding to the monomer exhibited the distinct colouration of F_420_ **(Supplementary Figure S1d)**. Both peaks showed identical migration by SDS-PAGE **(Supplementary Figure S1e)**. Furthermore, MSMEG_2027 is thermostable in excess of what is required for application in biological systems, with a *T*_M_^50^ of 58.9 ± 0.2 for the apoprotein and 64.5 ± 0.3 for the F_420_-bound state **(Supplementary Figure S2)**. Collectively, these results indicate that MSMEG_2027 exhibits the basic characteristics to develop a F_420_-dependent de-dimerization tool and its structural and thermodynamic properties were thus explored further.

### Structural analysis of the F_420_:MSMEG_2027 complex

In order to understand the role of the N-terminus in MSMEG_2027, we obtained a crystal structure of full-length monomeric F_420_:MSMEG_2027 complex at 1.67 Å (*P*2_1_2_1_2_1_ space group; **Figure 1a, Supplementary Table S1**). The F_420_:MSMEG_2027 complex crystallized with a single polypeptide chain in the asymmetric unit. F_420_ purified from recombinant *M. smegmatis* containing predominantly five to seven glutamate residues was used to generate the complex and electron density sufficient to build four glutamate residues into the model, as shown by a polder OMIT electron denisty map (**Figure 1a,c**) [28]. Consistent with previous molecular dynamics simulations, the *B*-factors of the atoms within the polyglutamate chain increase with distance from the deazaflavin ring, indicating increasing mobility in the distant regions of the cofactor (**Supplementary Figure S3**) [24]. The N-terminal helix is shown to be crucial to F_420_-binding as W11, V12 and Q15 form a pocket around the phenol ring of F_420_ which explains our observation that the NΔ26-MSMEG_2027 variant is inactive. Q15 forms a hydrogen bond with the 8-OH group of F_420_ stabilizing the phenolate form of F_420_ that is responsible for fluorescence [29].

**Figure 1.**
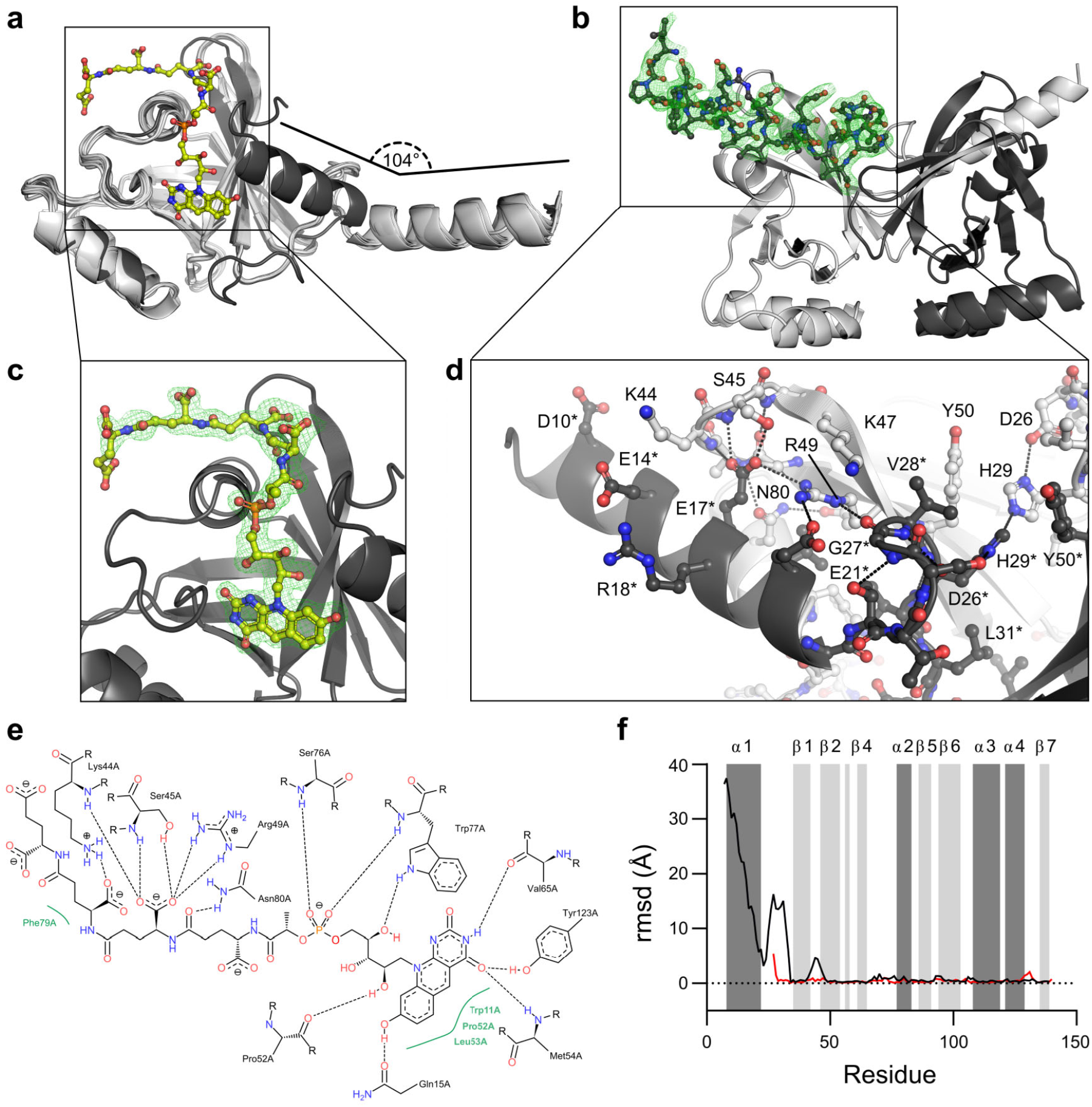
Structures of MSMEG_2027 in monomeric holo and dimeric apo forms. **a** Overlay of apoprotein chains (light grey) and the F_420_-bound structure (dark grey). In the apo complex, the N-terminal helix is orientated approximately 104° away from its position in the F_420_-bound structure, nearly orthogonal to the β-sheet. Additional short peptide in the F_420_-bound structure omitted for clarity. **b** Dimer composed of chains A and F with the position of the N-terminus shown by a polder omit map contoured at 3 Å. The orientation of the N-terminal helix results in a domain-swapped dimer in which the helix partially occupies the F_420_-binding site. **c** Polder omit map of F_420_ in the F_420_-bound structure contoured at 2 Å. Four glutamate residues of the F_420_ sidechain are clearly resolved. **d** Hydrogen bonding interactions between the N-terminal helix and the polyglutamate binding cleft in the dimer. **e** Ligand interaction diagram of F_420_-4 with MSMEG_2027. **f** Per-residue RMSD of Cα atoms between apoprotein chain F and the F_420_-bound structure (black line) and the NΔ26 truncated structure (red line). Regions forming α-helices (dark grey) and β-strands (light grey) are shaded.

### MSMEG_2027 forms a domain-swapped dimer in the absence of F_420_

To understand the nature of MSMEG_2027 dimerization in the absence of F_420_, we then solved the structure of the full-length dimeric apoenzyme at 2.37 Å (*P*2_1_ space group; **Supplementary Table S1**). In contrast to the small asymmetric unit of the F_420_-bound monomer, the asymmetric unit of this apoenzyme crystal contained 14 chains. Comparison between the full-length structures reveals that the N-terminus forms an α-helix in both cases (D10–Q22 in the F_420_-bound complex, T9–E21, in the apoenzyme, **Figure 1b**) but the orientation of this helix is divergent. In the F_420_-bound structure, the N-terminal helix lies in a perpendicular orientation over the central β-sheet, where it forms part of the F_420_ binding pocket around the deazaflavin moiety of F_420_ (**Figure 1c**), consistent with previously published structures of other FDOR-A proteins in complex with F_420_ [30]. In the apoenzyme structure, however, the helix engages in domain-swapping and is oriented almost perpendicular to the plane of the β-sheet, rotated 104° from the orientation in the F_420_-bound structure, and projects into the F_420_-binding pocket of the opposite subunit to interact with the T41–R49 loop (**Figure 1d**). This results in a parallel head-to-head dimer, as shown by a polder OMIT map of the N-terminal region (**Figure 1d**). The T41–R49 loop is displaced by up to 4 Å in the F_420_-bound monomer *vs*. the dimer. The displacement of this loop facilitates hydrogen bonding and the formation of salt bridges between K44, S45, and R49 with the second and third glutamate residues of F_420_. In contrast, these residues instead form hydrogen bonds or salt bridges with E17*, E21* and D26* in the dimeric apoenzyme structure. Additional hydrogen bonds from R13* to A43, S76 to L20* and a salt bridge between K44 to E14* are also observed.

The loop between the central β-sheet and N-terminal helix is also substantially reoriented with and RSMD greater than 5 Å for all mainchain atoms (aside from G23). In the dimer, this loop threads into the binding pocket of the deazaflavin core of F_420_ (**Figure 1b,d**) with V30 and L31 bridging the hydrophobic cores of the two chains. This conformation is stabilized by intrachain hydrogen bonding between E21 and G27, E21 and T24, as well as symmetrical interchain hydrogen bonds between D26 and H29* and D26* and H29 (**Figure 1b,d**). In the F_420_-bound structure, V30 and L31 instead form the substrate/cofactor binding pocket alongside W11, I35, M54 and Y120. The interaction between D26 and H29 observed in the dimer is disrupted in the F_420_-bound monomer and the sidechains of these residues are oriented away from each other. T24 and T25 form hydrogen bonds with G92 and D93, which comprise the turn between β−sheets β6 and β7. The interdomain distances in the domain swapped dimer are 58 and 40 Å at the N- and C-termini, respectively.

### Truncated MSMEG_2027 crystallizes in a pseudo-dimeric lattice

When the truncated apoenzyme structure is compared to the dimeric apoenzyme, the structures are almost identical in terms of a RMSD (**Figure 1f**). Interestingly, despite the absence of the N-terminus and the monomeric behaviour in solution, a pseudo-dimer of the truncated structure may be generated through application of the crystallographic two-fold symmetry operation (−*x, y*, −*z*) which is essentially identical to the full-length domain swapped dimer (RMSD 0.29 Å over 192 atoms, **Supplementary Figure S4**). Analysis of this pseudo-dimer with QtPISA [31] showed that the interface is much smaller in the pseudo-dimer than that of the full-length dimer (1026 Å^2^ *cf*. 3667–3867 Å^2^, representing 17% and ∼28% of the total solvent-accessible surface area, respectively) and that it lacks almost all hydrogen bonds and all salt bridges observed in the full-length dimer, as these are primarily associated with the domain-swapped N-terminal helix. No stable quaternary complex of the truncated protein was predicted by QtPISA, however the observation that the truncated protein aggregates into the crystal lattice in an identical orientation to how the full-length protein dimerises in solution suggests that dimerization is not solely dependent on the N-terminus and that the protein is predisposed to associate in this orientation. Similar pseudo-dimers are not observed in the crystal lattices of other truncated FDOR-A structures (PDB IDs 3R5L, 3R5P, 3R5R, and 3R5W) [30].

The recently reported structure of the full-length FDOR-A MSMEG_2850 (PDB ID 8D4W [32]), which shares 37% identity with MSMEG_2027, shows that it also forms a dimer in the absence of F_420_. However, in contrast to MSMEG_2027, dimerization of MSMEG_2850 occludes the cofactor binding site through a flexible loop region rather than domain-swapping of the N-terminus. The domain-swapped dimer of MSMEG_2027 also differs substantially from that seen in the other major class of mycobacterial FDORs, the FDOR-Bs. In the FDOR-B structures solved to-date the proteins form dimers, like MSMEG_2027, yet the dimer interface involves the opposite face of the β-sheet to that involved in cofactor binding, whereas in MSMEG_2027 the interface occurs on the same face as cofactor binding and occludes this binding site [23]. The structure of Rv1558:F_420_, an ortholog of MSMEG_2027 in *Mycobacterium tuberculosis* (49% amino acid identity to MSMEG_2027), shows a dimer topology similar to that seen in the FDOR-Bs [33]. Thus, the particular domain-swapped dimer architecture observed in MSMEG_2027 is not a widely conserved feature across all FDORs.

### Thermodynamic analysis of F_420_ binding and de-dimerization

To further characterise F_420_ binding and associated dissociation of MSMEG_2027, we performed isothermal titration calorimetry (ITC). Titrations of F_420_ into MSMEG_2027 produced strong and systematically increasing tailing of the thermogram peaks that necessitated increasing the injection interval to 600 s to ensure complete return to the baseline (**Figure 2a**). This is consistent with a (de)aggregation process associated with ligand binding [34]. The data could be fit with a one-site binding model in SEDPHAT [35] with an apparent *K*_D_ of 486 nM (68.3%; CI 378–611 nM; **Figure 2a**) in close agreement with results from intrinsic tryptophan fluorescence [23]. Fitting of the calorimetry data for titration of F_420_ into MSMEG_2027 to a one-site binding model captures the contributions of both F_420_ binding and dimer dissociation on the thermodynamic parameters. Binding of F_420_ to MSMEG_2027 was exothermic with a Δ*H* of −30.7 kcal mol^−1^ (68.3%; CI −32.0 to −29.5), −*T*Δ*S* of 22.1 kcal mol^−1^ (68.3%; CI 21.1 to 23.1) and a stoichiometry of 1:1 (**Figure 2b**). This substantial enthalpic component is consistent with the large number of electrostatic interactions formed between F_420_ and MSMEG_2027 in the crystal structure (**Figure 1**). There is an entropic cost to complex formation, despite dissociation of the dimer into monomers, most likely owing to the stabilisation of the otherwise flexible F_420_ polyglutamate chain and bound water molecules.

**Figure 2.**
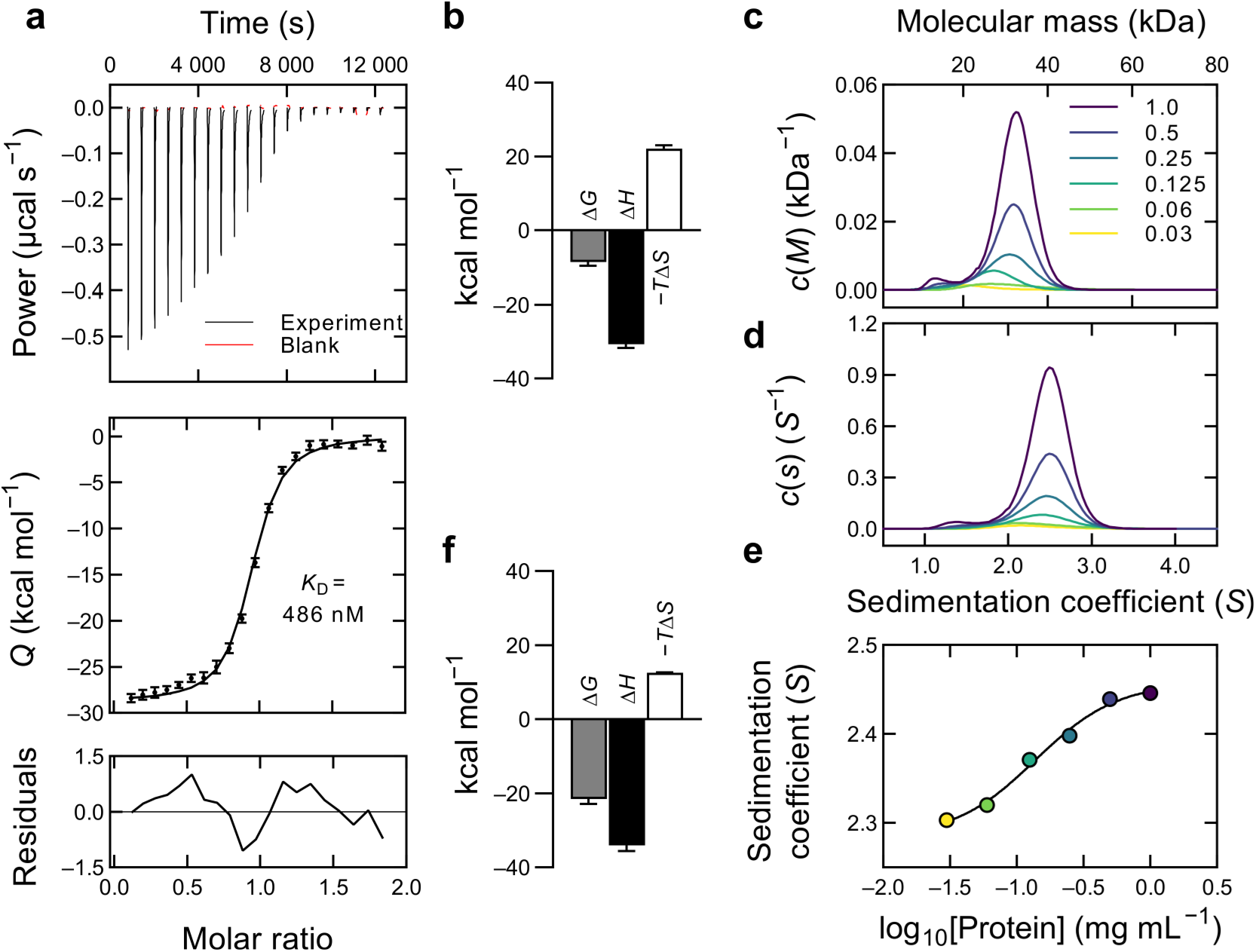
Biophysical characterisation of MSMEG_2027 interactions. **a** Representative titration of MSMEG_2027 (37.5 μM as monomer) with F_420_ (239 μM). Top panel shows the thermogram along with a control titration of F_420_ into buffer, middle panel shows the isotherm with the fit to a one-site binding model, bottom panel shows the residuals of the fit. The *K*_D_ was determined to be 486 nM (68.3% CI 378–611 nM). **b** Thermodynamic signature of F_420_ binding to MSMEG_2027. Error bars show the 68.3% CI of the calculated parameters. **c** Continuous mass sedimentation distribution of apo MSMEG_2027 as a function of molecular weight. Decreasing protein concentration results in collapse of discrete populations of monomer and dimer into a rapid equilibrium. Protein concentrations are in mg mL^−1^. **d** Continuous sedimentation distribution of MSMEG_2027 as a function of sedimentation coefficient. **e** The weight-average sedimentation coefficient from the integration of the distribution for each concentration is plotted as a function of protein concentration. The midpoint was determined to be 0.14 mg mL^−1^, corresponding to 8 μM on a monomer basis. **f** Thermodynamic signature of MSMEG_2027 dimerization obtained by analysis of the crystal structure with QtPISA. Note that the signs have been reversed to show the associative direction.

Having established that F_420_ binds with high (sub-micromolar) affinity and results in an exothermic dissociation of the dimer, we sought to investigate the affinity of the apoenzyme dimer in the absence of F_420_. To calculate the monomer-dimer equilibrium, we utilized analytical ultracentrifugation (AUC). Sedimentation velocity experiments at 1.0 mg mL^−1^ showed ∼96% of the protein exists in the dimeric form with a very small peak corresponding to ∼4% monomer (**Figure 2c**). Rather than a uniform shift in the relative peak heights with reduced protein concentration, we observed a broadening of the distribution at low concentrations suggesting that the protein enters into a rapid equilibrium between monomer at dimer rather than full dissociation into monomers [36] (**Figure 2c**). The weight-averaged sedimentation coefficient was obtained by integration analysis and the midpoint sedimentation coefficient determined to be 0.14 mg mL^−1^ (∼8 μM, **Figure 2d,e**).

We also investigated the dimer dissociation constant by ITC dilution experiments (**Supplementary Figure S5**) [37]. The isotherm clearly showed that the dimer dissociation corresponds to positive enthalpy, i.e., that protein dissociation is endothermic. This contrasts with the F_420_-dependent dissociation which is exothermic and enthalpically driven. The isotherm indicated a *K*_D_ on the order of 1.2 μM, although the quality of the isotherm was insufficient to derive an accurate value, potentially because of the rapid association and dissociation of monomers as indicated by the AUC data [38]. Computational analysis of the dimerization (using QtPISA) was consistent with the structural analysis, AUC and dilution ITC, indicating that dimerization is enthalpically driven (Δ*H* of −34.0 kcal mol^−1^ and −*T*Δ*S*^diss^ of 12.5 kcal mol^−1^, **Figure 2f**).

Collectively, several properties of MSMEG_2027 make it suitable for use in a chemogenetic F_420_-dependent de-dimerization system. First, it is a small (∼150 amino acids in length), thermostable protein that can be heterologously expressed in soluble form. Second, in the absence of F_420_, the protein forms a high affinity dimer that does not readily dissociate. Finally, F_420_ binds to the protein with high affinity (486 nM) and effectively induces dissociation of the dimer.

### Expression of MSMEG_2027 in human cells

Having established that the F_420_:MSMEG_2027 system meets several of the criteria for a CDD module, we set out to employ F_420_:MSMEG_2027 for the control of human cell signalling. We first evaluated whether MSMEG_2027 can be expressed in human cells. We fused a mammalian codon optimized MSMEG_2027 gene (see **Supplementary Information** for details) to the small fluorescent protein mVenus (mV) in a mammalian expression vector, followed by transfection into human embryonic kidney 293 (HEK293) cells. Fluorescence of mV is an established marker to quantify expression and visualise undesired aggregation [39, 40], and HEK293 cells represent a common cell model for signalling studies. Using bulk fluorescence measurements and fluorescence microscopy, we found that both N- and C-terminally fused MSMEG_2027 expressed efficiently and uniformly throughout the cytosol (**Figure 3a, Supplementary Figure S6**). Robust expression for both N-terminal and C-terminal fusions of MSMEG_2027 indicated independence of domain orientation. We did not observe aggregation and expression was only a few fold lower than the small highly expressing human FKBP protein.

**Figure 3.**
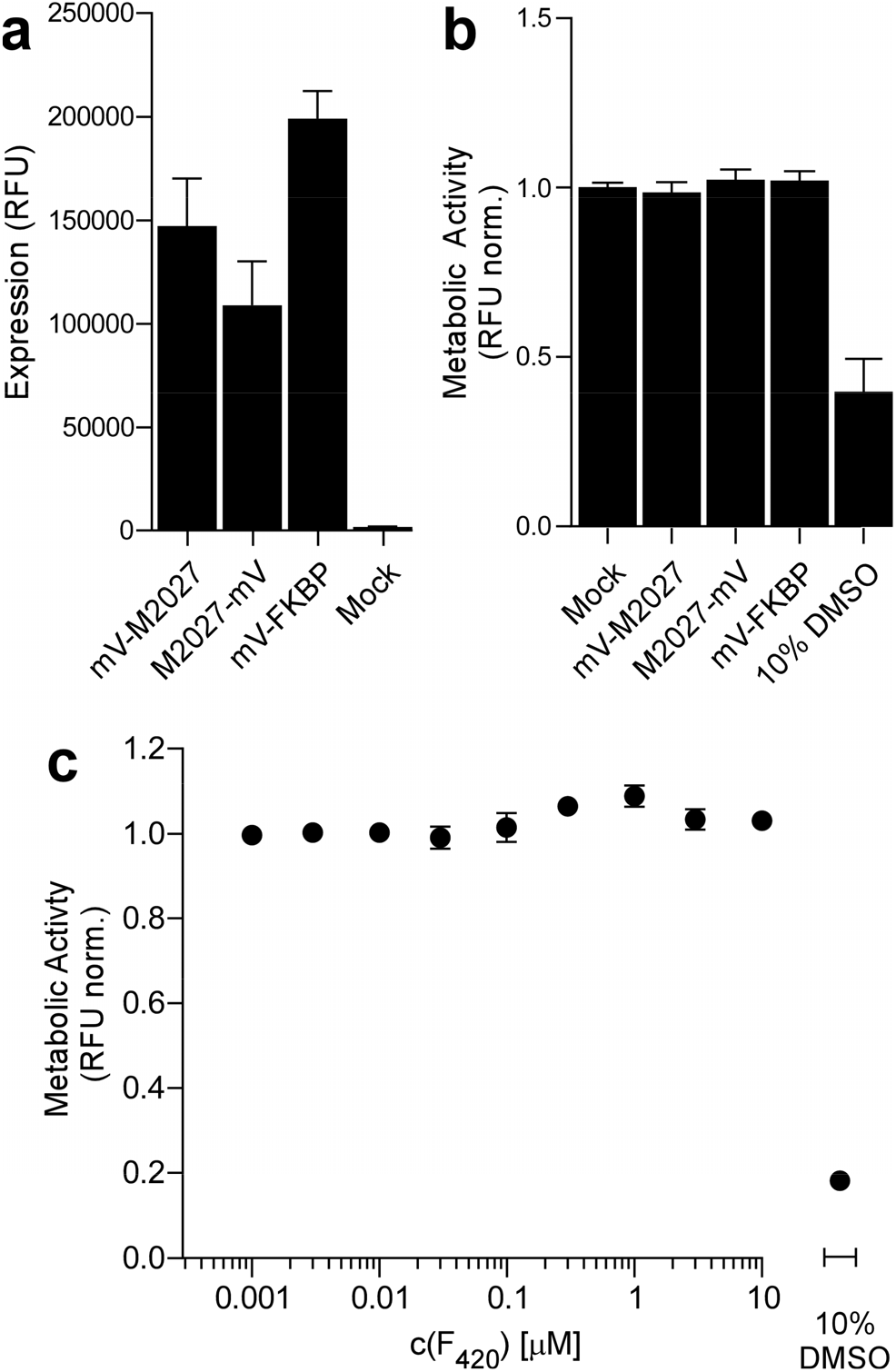
Expression of MSMEG_2027 and cell viability upon F_420_ treatment. **a** Expression measured as raw fluorescence units (RFU) for N- or C-terminally mV-tagged MSMEG_2027 (M2027) in HEK293 cells, compared to mV-tagged FKBP expressing and mock transfected cells. **b** Viability (measured as metabolic activity, normalized to mock transfected cells) of HEK293 cells expressing mV-tagged MSMEG_2027 and FKBP, compared to a 10% DMSO treated control with reduced viability. **c** Viability (measured as metabolic activity, normalized to vehicle treated cells) of HEK293 cells upon 24 h treatment with F_420_ compared to DMSO treated controls. In a–c, mean values ± SEM for three independent experiments, each performed in triplicate, are given.

We next validated that expression of MSMEG_2027 and application of F_420_ does not impact cell viability. Viability was assessed as metabolic activity through conversion of resazurin to resorufin as in our previous work on synthetic domain tools [7]. Cells transfected with mV-tagged MSMEG_2027 did not show reduced resorufin fluorescence compared to mV-tagged FKBP, suggesting that there is no negative effect of MSMEG_2027 expression on cell viability (**Figure 3b**). F_420_ is a 5-deazaflavin compound that is not found in eukaryotes. We verified that F_420_ does not exhibit toxicity in human cells in the concentration range required to induce de-dimerization of MSMEG_2027. We found that cell treatment with F_420_ for 24 h was well tolerated and did not impact metabolic activity and viability (compared to a vehicle treated control and across a concentration range of 0.001 to 10 μM) (**Figure 3c**). Thus, the chemical de-dimerizer F_420_ is tolerated at concentrations greater than the *K*_D_ for its binding to MSMEG_2027, suggesting that it should be possible to achieve essentially complete dimer dissociation without ligand-associated toxicity.

### MSMEG_2027-induced constitutive receptor activity is released by F_420_

Next, we demonstrated the applicability of F_420_:MSMEG_2027 as a CDD tool in an engineered receptor protein for chemically induced manipulation of cell signalling. The fibroblast growth factor receptor 1 (FGFR1) is a prototypical member of the large receptor tyrosine kinase (RTK) family that are major regulators of development and disease. Ligands of FGFRs (and other RTKs) induce receptor dimerization and FGFR1 can likewise be activated by forced dimerization [39, 41]. We engineered a chimeric receptor (MSMEG_2027-FGFR1) in which MSMEG_2027 replaces the extracellular domain of FGFR1 that normally binds FGF and induces dimerization, rendering the receptor insensitive to its natural ligand and solely under control of MSMEG_2027 (**Figure 4a**). We expected that attachment of MSMEG_2027 to FGFR1 would result in constitutive activation of the canonical MAPK/ERK signalling pathway that can then be disrupted by addition of F_420_ (**Figure 4a**). Indeed, HEK293 cells transfected with MSMEG_2027-FGFR1 showed MAPK/ERK pathway activity even to a similar extent as a constitutively active dimeric control receptor (FGFR1 fused to the dimeric IgG domain [40, 42]). In the next step, we tested whether this activity can be inhibited by addition of F_420_, and we found that signalling downstream of MSMEG_2027-FGFR1 was effectively reduced to levels comparable to a monomeric MSMEG_2027-deficient control receptor (**Figure 4b**).

**Figure 4.**
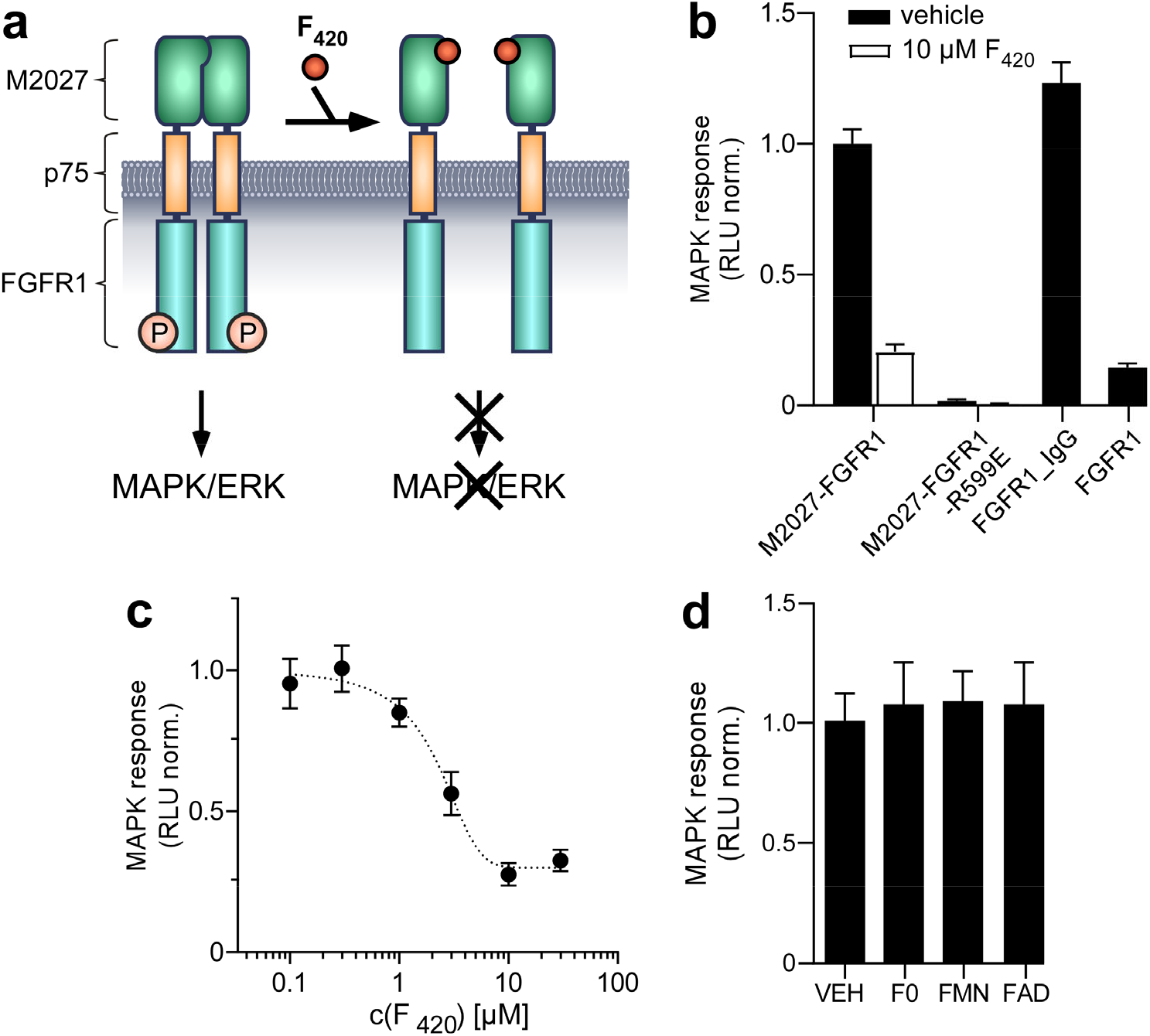
Regulation of cell signalling activity using MSMEG_2027 (M2027) and F_420_. **a** MSMEG_2027 was fused to FGFR1 to create a synthetic receptor that is constitutively active and inactivated by F_420_. **b** Activation of the MAPK/ERK pathway reporter expressed as raw luminescence units (RLU) by FGFR1 fused to MSMEG_2027, the Fc domain of IgG1 (IgG), or without modification, in response to treatment with 10 μM F_420_. Data are normalized to MSMEG_2027-FGFR1. **c** MAPK/ERK reporter response of MSMEG_2027-FGFR1 transfected cells to an increasing dose of F_420_, normalized to vehicle controls. **d** MAPK/ERK reporter response of MSMEG_2027-FGFR1 transfected cells to 10 μM F0, FMN and FAD. Data are normalized to vehicle controls. In b-d, mean values ± SEM for three to eleven independent experiments, each performed in triplicate, are given.

Motivated by the effective control of signalling, we examined the activation mechanism further. First, as a probe for dimer formation during receptor activation, we employed a charge inversion substitution (R599E numbered from the start of the construct, see **Materials and Methods**) in the FGFR1 kinase domain that prevents formation of a functionally essential, asymmetric dimer [43]. Indeed, MSMEG_2027-FGFR1-R599E did not induced MAPK/ERK pathway activation (**Figure 4b**) indicating that receptor dimerization through MSMEG_2027 is required. Second, we confirmed that MSMEG_2027-FGFR1 inactivation is dose-dependent. When testing a range of F_420_ concentrations on MSMEG_2027-FGFR1 expressing cells, 5 μM was sufficient to inhibit the MAPK signal to a basal level (**Figure 4c**). Finally, we confirmed that F_420_ binds to MSMEG_2027 with high specificity. F_420_ is chemically similar to FMN, flavin adenine dinucleotide (FAD), and F0, a chemical precursor of F_420_ consisting of just the core ring without phosphate, lactyl-group and polyglutamate chain [44-46]. We tested the response of MSMEG_2027-FGFR1 to these cofactors and found that neither F0, FMN nor FAD inhibited the MAPK/ERK pathway in receptor transfected cells (**Figure 4d**). Thus, MSMEG_2027 and F_420_ function orthogonally from these metabolites.

## DISCUSSION

The formation and disruption of PPIs is an abundant molecular mechanism underlying a plethora of cellular processes. Recent years have seen the rise of chemical and protein tools to manipulate PPIs *in situ* with methods for PPI induction being more abundant than those for disruption. Irreversible chemical disruption of PPIs was achieved using bifunctional crosslinkers that initially bridge common protein surface functional groups and are then cleaved either chemically or by light. The available chemogenetic protein-based disruption methods either rely on the competitive displacement of one binder by another or are limited to heteromeric interactions or large complexes. Thus, these also require the incorporation of several protein domains and dosing with several small molecule ligands in many cases.

Here, we have identified a novel, biorthogonal, and genetically encoded CDD tool and applied it for graded inhibition of cell signalling activity in human cells. We showed that MSMEG_2027, a small, thermostable mycobacterial protein, forms a unique domain-swapped homodimer, mediated primarily through its N-terminal helix. This dimer is disrupted by addition of F_420_, a cofactor of non-eukaryotic origin that is easy to produce and purify. We have found that binding affinities of F_420_ to MSMEG_2027 and of two MSMEG_2027 domains into the stoichiometrically defined dimer are in the high nanomolar and low micromolar range, respectively. MSMEG_2027 binds F_420_ with specificity over the ubiquitous flavin cofactors FMN and FAD. Signalling activity induced by MSMEG_2027-FGFR1 was unaffected by the addition of 10 μM FMN or FAD, which corresponds to >10-fold higher concentrations than the natural abundance of these flavins [47]. MSMEG_2027 expresses well in cells and both MSMEG_2027 and F_420_ were found to be non-toxic in a mammalian system. The new receptor MSMEG_2027-FGFR1 is, to our best knowledge, the first demonstration of ligand-based inhibition of a RTK where the ligand does not target an endogenous receptor domain (e.g., the kinase domain).

Our F_420_:MSMEG_2027 CDD module complements existing synthetic biology tools by providing a novel mechanism and chemistry independent from previously described domains, drugs and interaction modes. We are not aware of other proteins in which a bifurcational structure linking two domains is displaced by monomeric molecule causing dissociation. Expanding the synthetic biology toolbox with protein domains that recognize unique orthogonal cues is a prerequisite to delineate complex cellular processes, e.g., in tandem manipulation of processes driven by PPIs. The discovery of this minimal (i.e., one protein domain and one ligand) system highlights once more that Nature provides a repertoire of unique domains and multi-domain proteins.

## Supporting information

Supplemental Figures S1-S7, Tables S1-4, References

## Abbreviations

(CDD): Chemogenetic de-dimerization
(FGFR1): fibroblast growth factor receptor 1
(FDOR): flavin/deazaflavin oxidoreductase
(FAD): flavin adenine dinucleotide
(FMN): flavin mononucleotide
(FKBP): FK506 binding protein
(mV): mVenus
(ITC): isothermal titration calorimetry
(PPI): protein-protein interaction.

## ACKNOWLEDGEMENTS

We thank J. Kaczmarski for advice on isothermal titration calorimetry and helpful advice. This study was supported by grants of the Australian Research Council (FT200100519 and DP200102093, to H.J.; DE190100806 and DP220101901, to T.P.S.D.C; DP200102093, CE200100029, CE200100012 to C.J.J.), the National Health and Medical Research Council (APP1187638, to H.J.). S.K. was supported by the graduate program MolecularDrugTargets (Austrian Science Fund FWF), W1232). The Australian Regenerative Medicine Institute is supported by grants from the State Government of Victoria and the Australian Government. The EMBL Australia Partnership Laboratory (EMBL Australia) is supported by the National Collaborative Research Infrastructure Strategy (NCRIS) of the Australian Government. T.P.S.D.C. acknowledges the University of Adelaide for a Future Making Fellowship. E.R.R.M acknowledges the Grains Research and Development Corporation (9176977) for support through a PhD scholarship and operational funding. J.A. and E.R.R.M. were supported by the Australian Research Graduate Training Program scholarship. MicroMon of Monash University provided Sanger sequencing services. CJ thanks the ARC Centre of Excellence for Innovations in Peptide and Protein Science and the ARC Centre of Excellence in Synthetic Biology. We thank the staff of the MX2 beamline at the Australian Synchrotron, part of ANSTO, which made use of the Australian Cancer Research Foundation (ACRF) detector.

## AUTHOR CONTRIBUTIONS (CrediT taxonomy)

Conceptualization, S.K., C.J.J., H.J.; Funding Acquisition, T.P.S.D.C., C.J., H.J.; Methodology, J.A., S.K., E.R.R.M., T.P.S.D.C., C.J.J., H.J.; Project Administration, C.J.J., H.J.; Investigation, J.A., S.K., F.H.A., S.W.K., E.R.R.M., T.P.S.D.C.; Data curation, J.A., S.K., F.H.A., S.W.K., E.R.R.M., T.P.S.D.C.; Supervision, T.P.S.D.C., C.J., H.J.; Visualization, J.A., S.K., T.P.S.D.C., C.J.; Writing - Original Draft, J.A., S.K., E.R.R.M., T.P.S.D.C.; Writing - Review and Editing, C.J.J., H.J.

## MATERIALS AND METHODS

### Plasmid construction

Constructs of full-length and NΔ26 truncated MSMEG_2027 (UniProt ID A0QU01) in pETMCSIII for expression in *E. coli* were as described previously [23]. For mammalian expression a synthetic gene fragment encoding residues 2 to 140 of MSMEG_2027 was obtained with mammalian codon optimization (*H. sapiens*) following the manufacturer’s recommendation (Bioneer) (**Supplementary Table S2**) as for other gene sequences previously [48]. The fragment was amplified using PCR with overhanging restriction sites for *Cla*I (oligonucleotides 1 and 2, **Supplementary Table S3**) or *Age*I and *Xma*I (oligonucleotides 3 and 4, **Supplementary Table S3**) for incorporation into expression vectors.

For expression and viability testing, 2027 was inserted into the *Age*I restriction site of a previously described expression plasmid in pcDNA3.1(−) [39] that contains the fluorescent protein mV [49] preceded or followed by a glycine- and serine-rich linker and a compatible *Bsp*EI restriction site. A previously described mV-FKBP fusion protein was used as positive control [39].

To generate chimeric, de-dimerising receptors, MSMEG_2027 was cloned into a plasmid encoding the ligand binding and transmembrane domains of p75 (low affinity nerve growth factor receptor) and the kinase domain of murine FGFR1 in a pcDNA3.1(−) backbone (termed p75-FGFR1). MSMEG_2027 was inserted at a *Cla*I restriction site introduced between the signal peptide of p75 and the start of the extracellular domain through inverse PCR (oligonucleotides 5 and 6, **Supplementary Table S3**), yielding MSMEG_2027-FGFR1.

Point substitution R599E (numbered relative to the start codon of the fusion receptor) was introduced in MSMEG_2027-FGFR1 using site-directed mutagenesis PCR (oligonucleotides 7 and 8, **Supplementary Table S3**). The charge inversion substitution (R195E in full length murine FGFR1; R599E in our fusion receptor) prevents formation of a functionally essential, asymmetric kinase domain dimer in FGFR1 [43] and was used as a probe for dimer formation during receptor signalling as demonstrated previously [4, 42].

As a positive control construct, the Fc portion of human IgG1 (residues 230 to 461 of NCBI GenBank sequence KU951249.1 with an additional Cys to Ser substitution) was amplified from human cDNA using PCR using published protocols [50] (oligonucleotides 9 and 10, **Supplementary Table S3**) and inserted into p75-FGFR1 at an AgeI restriction site C-terminally of the FGFR1 kinase domain. The resulting construct yields a covalently linked receptor dimer that acts constitutively active [51]. All constructs were verified by DNA sequencing. Protein sequences are summarized in **Supplementary Table S4**.

### Production of F_420_

F_420_ was produced using a recombinant *M. smegmatis* system as described previously [52]. For cell-based experiments, F_420_ was dissolved in Tris-HCl adjusted to pH 7.5. Concentration was verified using a molar extinction coefficient of 41.4 mM^−1^ cm^−1^ at 420 nm [53]. For isothermal titration calorimetry experiments F_420_ was further purified on a HiLoad 16/60 Superdex 30 column (GE Healthcare) equilibrated with 0.3 M ammonium bicarbonate (pH 8.5). F_420_ containing fractions were pooled and evaporated to dryness. The residue was repeatedly redissolved in ultrapure water and evaporated to completely remove the ammonium bicarbonate and then dissolved in the same 1× phosphate buffered saline (PBS) used to prepare the receptor. Concentrations were determined using the pH-independent extinction coefficient of 25.7 mM^−1^ cm^−1^ at 400 nm [54].

### Protein expression and purification

The codon-optimized gene for full-length and NΔ26 truncated MSMEG_2027 in pETMCSIII were expressed and purified as described previously [23]. Briefly, the constructs were electroporated into *E. coli* BL21(DE3) cells and expressed in modified autoinduction media containing 2.0% tryptone, 0.5% yeast extract, 0.5% NaCl, 22 mM KH_2_PO_4_,42 mM Na_2_HPO_4_, 0.6% glycerol, 0.05% glucose, 0.2% lactose overnight at 30 °C. Cells were resuspended in lysis buffer (50 mM Na_2_HPO_4_, 300 mM NaCl, 25 mM imidazole, pH 8.0) and lysed by sonication on ice. The lysates were clarified by centrifugation at 20 000 × *g* for 1 hour at 4 °C before loading on a 5 mL HiTrap FF Ni-NTA column (GE Healthcare) equilibrated with lysis buffer. The column was washed with buffer containing 40 mM imidazole and target proteins eluted with 250 mM imidazole.

For crystallography the His_6_-tag was cleaved by overnight digestion with TEV protease expressed and purified in-house according to published protocols [55] in 50 mM Tris-HCl, 150 mM NaCl, 10 mM DTT, pH 8.0. Digestions were carried out at ambient temperature. Cleaved protein was separated from TEV protease by a subtractive Ni-NTA pass and loaded on to a HiLoad 16/60 Superdex 75 column (GE Healthcare) equilibrated with 20 mM Hepes, 150 mM NaCl, pH 7.5. In order to generate a holocomplex with F_420_ the concentrated protein sample was used to resuspend lyophilized F_420_ and incubated for 1 hour prior to size-exclusion. For binding studies size-exclusion was performed with 1× PBS (pH 7.4) supplemented with 1 mM CaCl_2_ and 0.5 mM MgCl_2_.

### Crystallization and structure determination

Fractions from size-exclusion corresponding to the monomer or dimer were exchanged into 20 mM Hepes, 50 mM NaCl, pH 7.5 and concentrated to 55 and 25 mg mL^−1^, respectively. Crystallization was performed using the hanging drop diffusion method at 18 °C with optimized conditions of 13% PEG 8000, 0.1 M Tris-HCl, 20% isopropanol for the dimer, and 0.07 M citrate, 0.03 M bis-tris propane, 16% PEG 3350 for the monomer. Crystal quality was improved with successive rounds of microseeding in both cases. Crystals were flash frozen in liquid nitrogen.

Data were collected at the Australian Synchrotron MX1 and MX2 beamlines for the dimer and monomer, respectively (wavelength 0.95373 Å). Data were processed and indexed with XDS [56] and scaled using AIMLESS [57] from the CCP4 suite [58] and are shown in **Table 1**. Both structures were solved by molecular replacement using PHASER [59] with the truncated MSMEG_2027 structure (PDB ID 4Y9I) as the search model. Parameters for F_420_-4 were generated with ACEDRG [60] and eLBOW [61]. Models were improved by manual model building with COOT [62] interspersed with refinement using REFMAC5 [63] and phenix.refine [64]. Ramachandran statistics for the final models were obtained using MOLPROBITY [65] and are as follows: Monomer 6WTA, 98% favoured, 2% allowed, 0% outliers; Dimer 6XRI, 97% favoured, 3% allowed, 0% outliers. Polder maps were generated using phenix.polder [28].

### Analytical ultracentrifugation

Sedimentation velocity experiments were performed in a XL-A analytical ultracentrifuge (Beckman Coulter) using an 8-hole An50-Ti rotor and double-sector cells containing synthetic quartz windows as previously described [66-68]. Cells were loaded with 380 μL of protein diluted in 1× PBS (pH 7.4) supplemented with 1 mM CaCl_2_ and 0.5 mM MgCl_2_ and 400 μL of reference solution. The samples were then centrifuged at 40 000 rpm at 25 °C. Data was collected at 280 nm in continuous mode without averaging using a step size of 0.003 cm. Solvent density (1.0056 g mL^−1^), buffer viscosity (0.0101987 cp) and the estimated sample partial specific volume (0.734516 mL g^−1^) were computed using SEDNTERP [36]. The continuous sedimentation [*c*(*s*)] and mass [*c*(*M*)] distributions were determined by fitting absorbance as a function of radial position to the Lamm equation using SEDFIT [69]. The weight-average sedimentation coefficients were determined by integration of the continuous sedimentation distributions using SEDFIT [69]. The weight-average sedimentation coefficients were plotted as a function of protein concentration for estimation of the dissociation constant (*K*_D_) [70].

### Isothermal titration calorimetry

Titrations were performed with a low volume Nano ITC calorimeter (TA Instruments) at 298.15 K with 400 rpm stirring. Titrations of F_420_ into MSMEG_2027 were carried out with 37.5 μM MSMEG_2027 (monomer basis) in the cell and 239 μM F_420_ in the syringe. The injection program delivered 1 μL of titrant for the first injection and 2 μL titrant for the subsequent 20 injections. Data were collected for 300 s prior to the first injection and for 600 s following each injection. Baselines were subtracted and heats integrated using NITPIC [71] and the data fit to a one-site binding model in SEDPHAT [35]. Experiments were repeated three times. Model fitting and global error estimation was performed based on existing protocols [34]. For dimer dissolution experiments MSMEG_2027 (1–2.9 mM) was titrated into buffer following the injection program outlined above except that injections were spaced at 350 s and the cell stirred at 350 rpm.

### Circular dichroism

Spectra were acquired with a ChiraScan spectrophotometer (Applied Photophysics) equipped with a Quantum temperature controller (NorthWest). Samples containing 11.25 μM MSMEG_2027 and 0–112.5 μM F_420_ were prepared in 10 mM sodium phosphate pH 7.5 and measured in a 1 mm QS quartz cuvette (Hellma Analytics). Spectra were recorded between 190 and 260 nm with a 1 nm slit width. For thermostability determinations spectra were recorded at 218 nm and the temperature increased from 20 to 90 °C at 1 °C min^−1^. *T*_M_^50^ was calculated by fitting to the Boltzmann sigmoid model in GraphPad Prism v8.4 (GraphPad Sofware Inc.).

### Mammalian cell culture and transfection

HEK293 cells (ThermoFisher; further authenticated by assessing cell morphology and growth rate) were maintained in full medium (DMEM supplemented with 10% FBS, 100 U mL^−1^ penicillin and 0.1 mg mL^−1^ streptomycin) at 37 °C in a humidified incubator with 5% CO_2_. All transfections were carried out in transfection medium (DMEM supplemented with 5% FBS) using polyethyleneimine (PEI) as previously described [40]. Experiments were conducted in 96-well clear bottom plates coated with poly-L-ornithine.

### Fluorescent fusion proteins, expression, and resazurin viability testing

For F_420_ toxicity testing, 5 × 10^4^ HEK293 cells were seeded per well and after 6 h treated with medium supplemented with 10% vehicle or F_420_ at the indicated concentrations. A positive control sample was treated with 10% DMSO to induce cell death. For expression and domain toxicity testing, 5 × 10^4^ HEK293 cells were transfected with 100 ng expression plasmid encoding mV fusions. Expression was assessed in a microplate reader 30 h after transfection. Viability was assessed as metabolic activity using resazurin [72]. Cells were incubated for 1 h with resazurin (0.01 mg mL^−1^), and fluorescence of the metabolic product resorufin was quantified in a microplate reader. Fluorescence microscopy images were recorded on a digital microscope (Nikon Eclipse Ti-S) 30 h after transfection.

### MAPK/ERK pathway activation

Activation of the MAPK/ERK pathway was assessed with the PathDetect Elk1 *trans*-Reporting System (Agilent) containing firefly luciferase. In brief, 5 × 10^4^ HEK293 cells per well were transfected with 213 ng DNA (200 ng *trans*-activator, 10 ng *trans*-reporter, 3 ng receptor) and 1000 ng PEI per well. After 6 h transfection medium was changed to starve medium (0.5% FBS) with 10% vehicle (Tris-HCl, pH 7.5), F_420_ (concentration as indicated) or 10 μM FMN, FAD or F0 (a metabolic precursor of F_420_). Cells were incubated for another 16 h, and luciferase expression was assessed with a homemade dual-luciferase assay reagent [73]. Luminescence was measured in a microplate reader.

### Accession numbers

Coordinates and structure factors of monomeric F_420_:MSMEG_2027 and dimeric apo MSMEG_2027 have been deposited in the PDB with accession numbers 6WTA (monomer) and 6XRI (dimer).

