## Supplemental Figures S1-S7, Tables S1-4, References for "A F_420_-dependent single domain chemogenetic tool for protein de-dimerization"

**SUPPLEMENTARY INFORMATION**

Antoney *et al.*

**Supplementary Information:**
Supplementary Figure S1-S7
Supplementary Tables S1-S4
Supplementary References

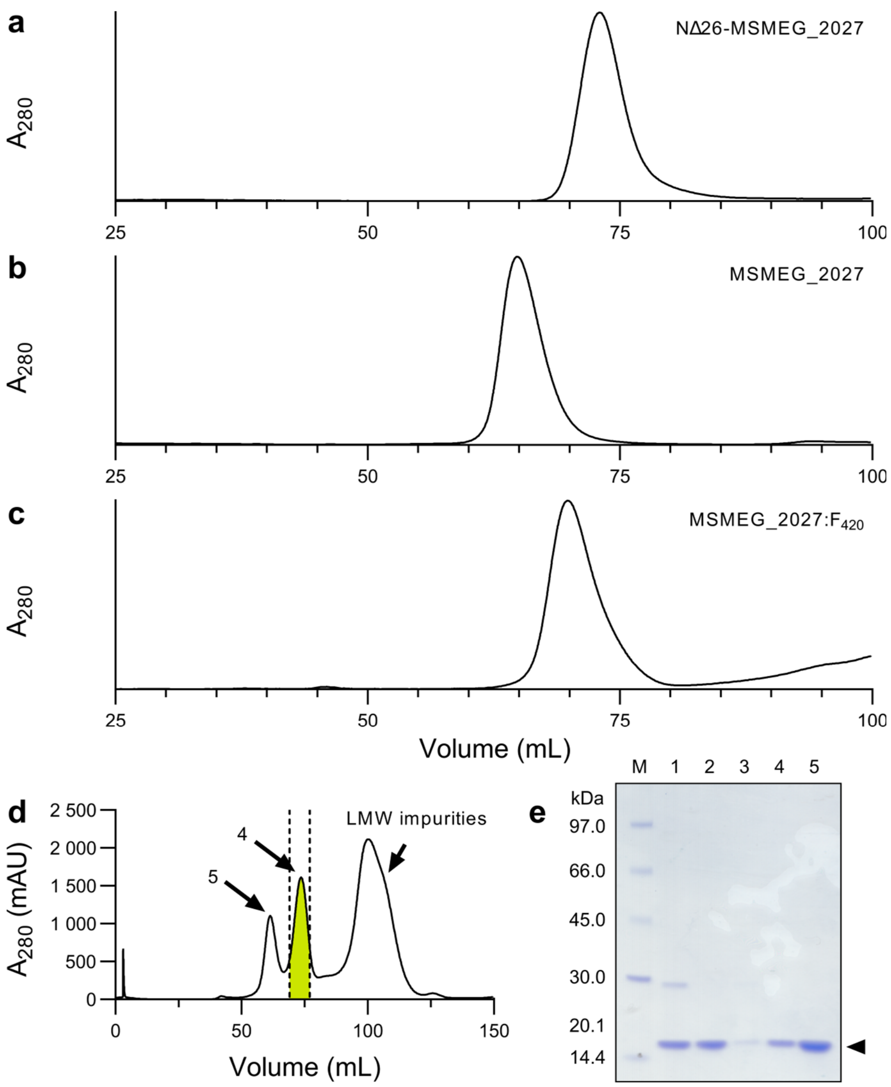


**Figure S1.** Gel filtration of MSMEG_2027 variants. **a** The NΔ26 truncated variant (MW = 14.0 kDa) eluted with $R_v$ = 73 mL, consistent with a monomer. **b** The full-length apoprotein (MW = 17.9 kDa monomer, 35.9 kDa dimer) elutes with $R_v$ = 65 mL, consistent with a dimer. **c** Complex of full-length protein with $F_{420}$ (MW = 19.2 kDa, assuming $F_{420}$-6 bound) elutes with $R_v$ = 70 mL. All peak heights were normalised for comparison. Samples were loaded on a HiLoad 16/60 Superdex 75 pg column (GE Healthcare) equilibrated with 20 mM Hepes, 150 mM NaCl, pH 7.5 and contained 10 to 45 mg of protein. **d** Preparative gel filtration of MSMEG_2027:$F_{420}$ with incomplete conversion to holocomplex. **e** SDS-PAGE analysis of purification in panel **d**. Lanes are 1: TEV digestion prior to subtractive Ni-NTA; 2: Flow through of subtractive Ni-NTA; 3: Eluate of Ni-NTA with 250 mM imidazole; 4: Second gel filtration peak with green colouration; 5: First colourless gel filtration peak. Note that $F_{420}$ absorbs strongly at 280 nm but is not retained following SDS-PAGE. The band corresponding to MSMEG_2027 is indicated with the marker.

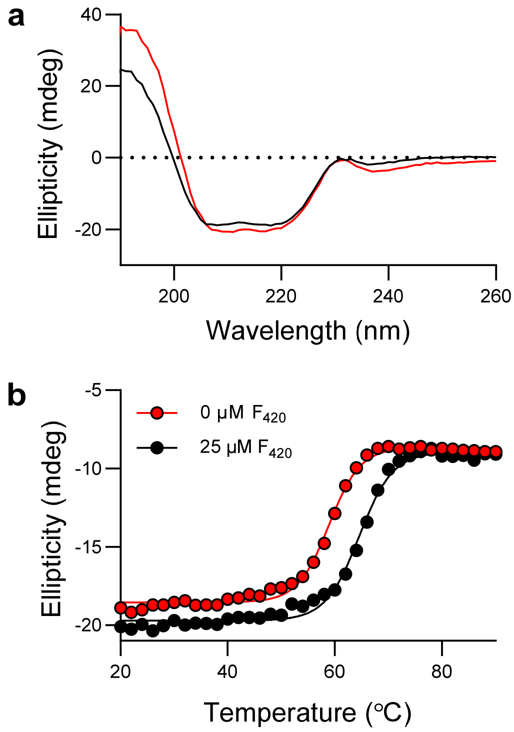


**Figure S2.** Thermal stability of MSMEG_2027. **a** Far-UV circular dichroism spectra of 11.3 µM MSMEG_2027 at 20°C with 0 (red line) and 28.1 µM $F_{420}$ (black line). All samples were prepared in 10 mM sodium phosphate (pH 7.5). **b** Thermal denaturation of MSMEG_2027 followed at 218 nm in the presence of varying concentrations of $F_{420}$. A temperature ramp of 1°C $min^{-1}$ was applied from 20 to 90°C. $T_M^{50}$ values were calculated by fitting the Boltzmann sigmoidal model in GraphPad Prism v8.4.

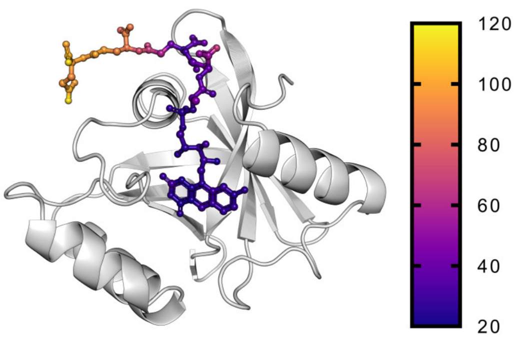


**Figure S3.** *B*-factors of $F_{420}$-4. Atoms of $F_{420}$-4 are coloured by *B*-factors. Consistent with molecular dynamics simulations [1], the third and fourth glutamate residues are more flexible that the first and second. The average *B*-factor for $F_{420}$-4 is 51.0 Å$^2$, while the factor of the protein is 48.0 Å$^2$.

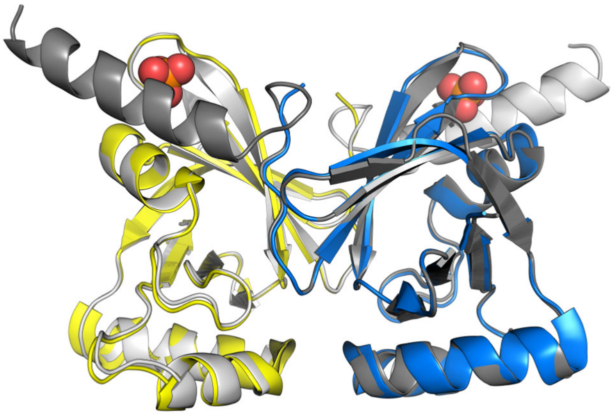

**Figure S4.** Pseudo-dimer of NΔ26 MSMEG_2027. Application of the twofold crystallographic symmetry operator (−*x*, *y*, −*z*) to the truncated structure (PDB: 4Y9I, yellow and blue) generates a pseudo-dimer of full-length MSMEG_2027 (white and grey). The structures superimpose with a RMSD of 0.29 Å over 192 atoms.

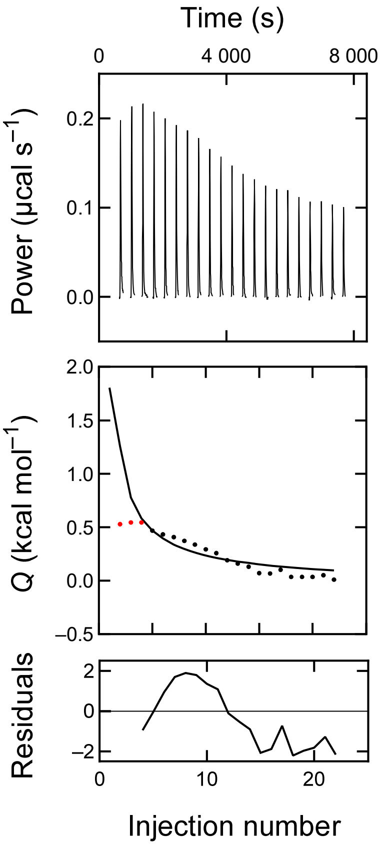


**Figure S5.** MSMEG_2027 dimer dilution titration. MSMEG_2027 (2.94 mM monomer basis) was titrated into 1× PBS (pH 7.4) supplemented with 1 mM $CaCl_2$ and 0.5 mM $MgCl_2$. Dimer dissociation was endothermic as indicated by the positive peak direction. Peak picking was done with NITPIC and the data reimported into NanoAnalyze. The constant heat obtained with the final injections was subtracted as the baseline and the model fit to the dimer dissociation model in NanoAnalyze, omitting the first three injections as they deviated substantially from the model. Although a best fit value for the $K_D$ in the low micromolar range was obtained, the poor fit to the model resulted in large error estimates spanning two orders of magnitude, and thus does not provide a reliable estimate of the $K_D$.

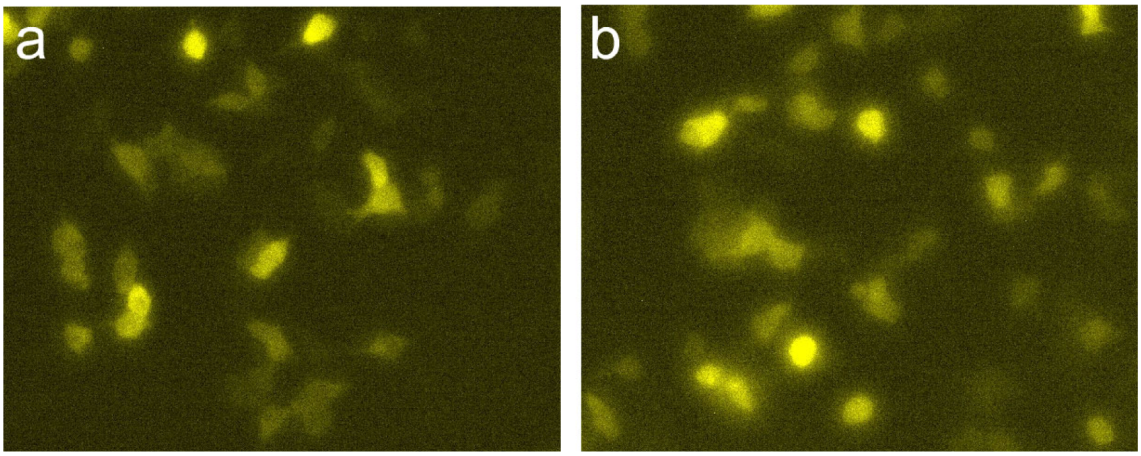


**Figure S6.** Expression of MSMEG_2027 in HEK293 cells. Representative fluorescence microscopy images of mVenus (mV) fusion proteins. **a** mV-MSMEG_2027 and **b** MSMEG_2027-mV in HEK293 cells. Images were taken in a Nikon Eclipse Ti-S microscope 30 h after transfection of cells.

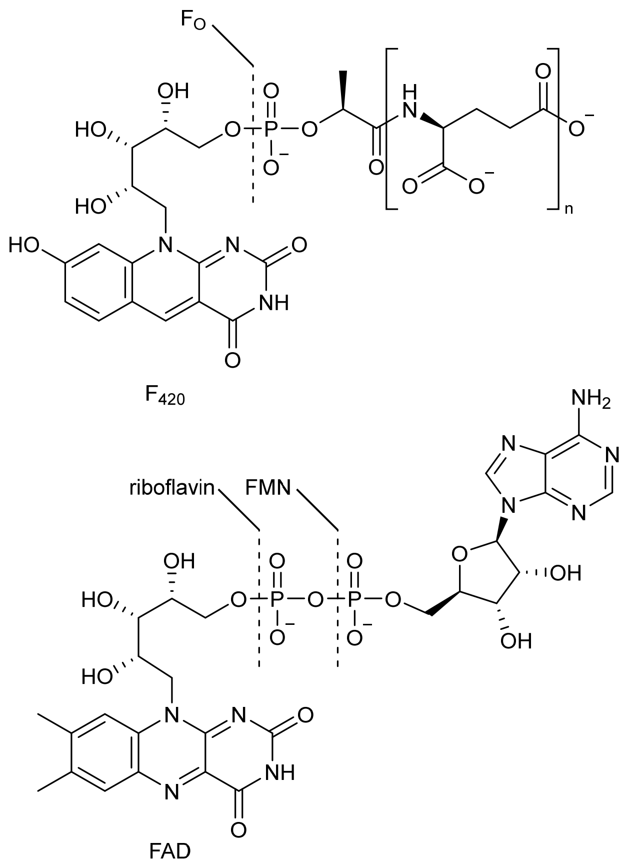


**Figure S7.** Structures of $F_{420}$, F0, FMN and FAD. $F_{420}$ is a derivative of 7,8-didemethyl-8-hydroxy-5-deazariboflavin, also called F0, containing between two and seven glutamate residues in the side chain.

**Table S1.** Data collection, processing and refinement statistics.

| | MSMEG_2027:$F_{420}$[a] PDB 6WTA | MSMEG_2027 dimer[a] PDB 6XRI |
|---|---|---|
| **Data collection** | | |
| Space group | $P2_12_12_1$ | $P2_1$ |
| Cell dimensions | | |
| *a*, *b*, *c* (Å) | 35.364, 63.022, 65.366 | 73.55, 110.23, 140.15 |
| $\alpha, \beta, \gamma$ (°) | 90, 90, 90 | 90, 105.21, 90 |
| Resolution (Å) | 32.68–1.67 (1.70–1.67)* | 34.21–2.37 (2.42–2.37)* |
| $R_{merge}$ | 0.173 (1.556) | 0.205 (1.703) |
| $R_{pim}$ | 0.057 (0.511) | 0.123 (1.007) |
| $I / \sigma I$ | 10.1 (1.9) | 3.9 (0.6) |
| $CC_{1/2}$ | 0.996 (0.910) | 0.994 (0.520) |
| Completeness (%) | 99.8 (99.9) | 92.9 (98.3) |
| Redundancy | 10.6 (10.1) | 3.7 (3.8) |
| **Refinement** | | |
| Resolution (Å) | 32.68–1.67 | 34.184–2.37 |
| No. reflections | 17596 (886) | 81254 (8565) |
| $R_{work}$ / $R_{free}$ | 20.0/24.1 | 24.92/30.41 |
| No. atoms | | |
| Protein | 1179 | 15316 |
| Ligand/ion | 71 | 0 |
| Water | 87 | 817 |
| *B*-factors | | |
| Protein | 48.02 | 48.8 |
| $F_{420}$-4 (n=1) | 51.03 | – |
| Water | 45.95 | 30.8 |
| R.M.S. deviations | | |
| Bond lengths (Å) | 0.004 | 0.0064 |
| Bond angles (°) | 0.69 | 0.70 |

*Values within parentheses are for highest-resolution shell.
[a] Number of crystals = 1.

**Table S2.** Mammalian codon-optimised sequence of MSMEG_2027.

| Sequence |
|---|
| ACAGATGCGGAACTTTCCCCTACTGACTGGGTGCGCGAGCAGACGGAGAGAATCCTTGAACAA<br>GGCACAACTGATGGGGTGCATGTCCTTGATCGGCCCATAGTACTTTTCACGACGACGGGAGCCA<br>AAAGCGGTAAGAAACGCTATGTGCCTTTGATGCGGGTAGAGGAGAACGGCAAGTACGCTATGGT<br>CGCTTCTAAGGGAGGCGACCCCAAGCACCCCTCCTGGTACTTCAATGTTAAAGCTAACCCTACA<br>GTGTCCGTTCAGGATGGGGACAAAGTTCTCCCCGATAGGACTGCCCGAGAGTTGGAAGGCGAG<br>GAGAGAGAGCACTGGTGGAAGTTGGCTGTAGAGGCGTACCCTCCCTATGCTGAATATCAAACCA<br>AAACTGACCGCCTGATTCCAGTTTTCATAGTTGAG |

**Table S3.** Oligonucleotides used in this study. Restriction sites or mutated bases are underlined.

| No. | Sequence |
|---|---|
| 1 | GATCATATCGATACAGATGCGGAACTTTCCCCTACTGACT |
| 2 | GATCATATCGATCTCAACTATGAAAACTGGAATCAGGCGGTCAGT |
| 3 | GATCATACCGGTACAGATGCGGAACTTTCCCCTACTGACT |
| 4 | GATCATCCCGGGTCAACTATGAAAACTGGAATCAGGCGGTCAGT |
| 5 | GATCATATCGATAAAGAAGCATGCCCAACTGGACTCTACAC |
| 6 | GATCATATCGATTGCTCCACCCAGGGACACT |
| 7 | GTATCTACAGGCCCGGGAGCCTCCTGGGCTGGAG |
| 8 | CTCCAGCCCAGGAGGCTCCCGGGCCTGTAGATAC |
| 9 | GATCATACCGGTGAGCCCAAATCTTCTGACAAAACTCACACAT |
| 10 | GATCATACCGGTTTTACCCGGAGACAGGGAGAGG |

**Table S4.** Protein sequences of full-length MSMEG_2027 and fusion proteins.

| Name | Sequence |
|---|---|
| MSMEG_2027 (UniProt: A0QU01) | TDAELSPTDWVREQTERILEQGTTDGVHVLDRPIVLFTTTGAKSGKKRYVPL MRVEENGKYAMVASKGGDPKHPSWYFNVKANPTVSVQDGDKVLPDRTARE LEGEEREHWWKLAVEAYPPYAEYQTKTDRLIPVFIVE |
| MSMEG_2027-mV | MSGTDAELSPTDWVREQTERILEQGTTDGVHVLDRPIVLFTTTGAKSGKKRY VPLMRVEENGKYAMVASKGGDPKHPSWYFNVKANPTVSVQDGDKVLPDRT ARELEGEEREHWWKLAVEAYPPYAEYQTKTDRLIPVFIVEPGGSSGSVSKGE ELFTGVVPILVELDGDVNGHKFSVSGEGEGDATYGKLTLKLICTTGKLPVPWP TLVTTLGYGLQCFARYPDHMKQHDFFKSAMPEGYVQERTIFFKDDGNYKTR AEVKFEGDTLVNRIELKGIDFKEDGNILGHKLEYNYNSHNVYITADKQKNGIKA NFKIRHNIEDGGVQLADHYQQNTPIGDGPVLLPDNHYLSYQSKLSKDPNEKR DHMVLLEFVTAAGITLGMDELYK |
| mV-MSMEG_2027 | MVSKGEELFTGVVPILVELDGDVNGHKFSVSGEGEGDATYGKLTLKLICTTG KLPVPWPTLVTTLGYGLQCFARYPDHMKQHDFFKSAMPEGYVQERTIFFKD DGNYKTRAEVKFEGDTLVNRIELKGIDFKEDGNILGHKLEYNYNSHNVYITAD KQKNGIKANFKIRHNIEDGGVQLADHYQQNTPIGDGPVLLPDNHYLSYQSKL SKDPNEKRDHMVLLEFVTAAGITLGMDELYKGSSGSSGTDAELSPTDWVRE QTERILEQGTTDGVHVLDRPIVLFTTTGAKSGKKRYVPLMRVEENGKYAMVA SKGGDPKHPSWYFNVKANPTVSVQDGDKVLPDRTARELEGEEREHWWKLA VEAYPPYAEYQTKTDRLIPVFIVEPG |
| FGFR1 | MGAGATGRAMDGPRLLLLLLLGVSLGGAIDKEACPTGLYTHSGECCKACNL GEGVAQPCGANQTVCEPCLDSVTFSDVVSATEPCKPCTECVGLQSMSAPC VEADDAVCRCAYGYYQDETTGRCEACRVCEAGSGLVFSCQDKQNTVCEEC PDGTYSDEANHVDPCLPCTVCEDTERQLRECTRWADAECEEIPGRWITRST PPEGSDSTAPSTQEPEAPPEQDLIASTVAGVVTTVMGSSQPVVTRGTTDNLI PVYCSILAAVVVGLVAYIAFKRGRAMKSGTKKSDFHSQMAVHKLAKSIPLRRQ VTVSADSSASMNSGVLLVRPSRLSSSGTPMLAGVSEYELPEDPRWELPRDR LVLGKPLGEGCFGQVVLAEAIGLDKDKPNRVTKVAVKMLKSDATEKDLSDLIS EMEMMKMIGKHKNIINLLGACTQDGPLYVIVEYASKGNLREYLQARRPPGLE YCYNPSHNPEEQLSSKDLVSCAYQVARGMEYLASKKCIHRDLAARNVLVTE DNVMKIADFGLARDIHHIDYYKKTTNGRLPVKWMAPEALFDRIYTHQSDVWS FGVLLWEIFTLGGSPYPGVPVEELFKLLKEGHRMDKPSNCTNELYMMMRDC WHAVPSQRPTFKQLVEDLDRIVALTSNQEYLDLSIPLDQYSPSFPDTRSSTCS SGEDSVFSHEPLPEEPCLPRHPTQLANSGLKRRVETGGSGVDYPYDVPDYA LD |
| MSMEG_2027-FGFR1 | MGAGATGRAMDGPRLLLLLLLGVSLGGAIDTDAELSPTDWVREQTERILEQG TTDGVHVLDRPIVLFTTTGAKSGKKRYVPLMRVEENGKYAMVASKGGDPKH PSWYFNVKANPTVSVQDGDKVLPDRTARELEGEEREHWWKLAVEAYPPYA EYQTKTDRLIPVFIVEIDKEACPTGLYTHSGECCKACNLGEGVAQPCGANQT VCEPCLDSVTFSDVVSATEPCKPCTECVGLQSMSAPCVEADDAVCRCAYGY YQDETTGRCEACRVCEAGSGLVFSCQDKQNTVCEECPDGTYSDEANHVDP CLPCTVCEDTERQLRECTRWADAECEEIPGRWITRSTPPEGSDSTAPSTQE PEAPPEQDLIASTVAGVVTTVMGSSQPVVTRGTTDNLIPVYCSILAAVVVGLV AYIAFKRGRAMKSGTKKSDFHSQMAVHKLAKSIPLRRQVTVSADSSASMNS GVLLVRPSRLSSSGTPMLAGVSEYELPEDPRWELPRDRLVLGKPLGEGCFG QVVLAEAIGLDKDKPNRVTKVAVKMLKSDATEKDLSDLISEMEMMKMIGKHK NIINLLGACTQDGPLYVIVEYASKGNLREYLQARRPPGLEYCYNPSHNPEEQL SSKDLVSCAYQVARGMEYLASKKCIHRDLAARNVLVTEDNVMKIADFGLARD IHHIDYYKKTTNGRLPVKWMAPEALFDRIYTHQSDVWSFGVLLWEIFTLGGSP YPGVPVEELFKLLKEGHRMDKPSNCTNELYMMMRDCWHAVPSQRPTFKQL VEDLDRIVALTSNQEYLDLSIPLDQYSPSFPDTRSSTCSSGEDSVFSHEPLPE EPCLPRHPTQLANSGLKRRVETGGSGVDYPYDVPDYALD |
| MSMEG_2027-FGFR1-R599E | MGAGATGRAMDGPRLLLLLLLGVSLGGAIDTDAELSPTDWVREQTERILEQG TTDGVHVLDRPIVLFTTTGAKSGKKRYVPLMRVEENGKYAMVASKGGDPKH PSWYFNVKANPTVSVQDGDKVLPDRTARELEGEEREHWWKLAVEAYPPYA EYQTKTDRLIPVFIVEIDKEACPTGLYTHSGECCKACNLGEGVAQPCGANQT VCEPCLDSVTFSDVVSATEPCKPCTECVGLQSMSAPCVEADDAVCRCAYGY |

| | |
|---|---|
| | YQDETTGRCEACRVCEAGSGLVFSCQDKQNTVCEECPDGTYSDEANHVDP<br>CLPCTVCEDTERQLRECTRWADAECEEIPGRWITRSTPPEGSDSTAPSTQE<br>PEAPPEQDLIASTVAGVVTTVMGSSQPVVTRGTTDNLIPVYCSILAAVVVGLV<br>AYIAFKRGRAMKSGTKKSDFHSQMAVHKLAKSIPLRRQVTVSADSSASMNS<br>GVLLVRPSRLSSSGTPMLAGVSEYELPEDPRWELPRDRLVLGKPLGEGCFG<br>QVVLAEAIGLDKDKPNRVTKVAVKMLKSDATEKDLSDLISEMEMMKMIGKHK<br>NIINLLGACTQDGPLYVIVEYASKGNLREYLQAREPPGLEYCYNPSHNPEEQL<br>SSKDLVSCAYQVARGMEYLASKKCIHRDLAARNVLVTEDNVMKIADFGLARD<br>IHHIDYYKKTTNGRLPVKWMAPEALFDRIYTHQSDVWSFGVLLWEIFTLGGSP<br>YPGVPVEELFKLLKEGHRMDKPSNCTNELYMMMRDCWHAVPSQRPTFKQL<br>VEDLDRIVALTSNQEYLDLSIPLDQYSPSFPDTRSSTCSSGEDSVFSHEPLPE<br>EPCLPRHPTQLANSGLKRRVETGGSGVDYPYDVPDYALD |
| FGFR1-IgG | MGAGATGRAMDGPRLLLLLLLGVSLGGAIDKEACPTGLYTHSGECCKACNL<br>GEGVAQPCGANQTVCEPCLDSVTFSDVVSATEPCKPCTECVGLQSMSAPC<br>VEADDAVCRCAYGYYQDETTGRCEACRVCEAGSGLVFSCQDKQNTVCEEC<br>PDGTYSDEANHVDPCLPCTVCEDTERQLRECTRWADAECEEIPGRWITRST<br>PPEGSDSTAPSTQEPEAPPEQDLIASTVAGVVTTVMGSSQPVVTRGTTDNLI<br>PVYCSILAAVVVGLVAYIAFKRGRAMKSGTKKSDFHSQMAVHKLAKSIPLRRQ<br>VTVSADSSASMNSGVLLVRPSRLSSSGTPMLAGVSEYELPEDPRWELPRDR<br>LVLGKPLGEGCFGQVVLAEAIGLDKDKPNRVTKVAVKMLKSDATEKDLSDLIS<br>EMEMMKMIGKHKNIINLLGACTQDGPLYVIVEYASKGNLREYLQARRPPGLE<br>YCYNPSHNPEEQLSSKDLVSCAYQVARGMEYLASKKCIHRDLAARNVLVTE<br>DNVMKIADFGLARDIHHIDYYKKTTNGRLPVKWMAPEALFDRIYTHQSDVWS<br>FGVLLWEIFTLGGSPYPGVPVEELFKLLKEGHRMDKPSNCTNELYMMMRDC<br>WHAVPSQRPTFKQLVEDLDRIVALTSNQEYLDLSIPLDQYSPSFPDTRSSTCS<br>SGEDSVFSHEPLPEEPCLPRHPTQLANSGLKRRVETGEPKSSDKTHTCPPC<br>PAPELLGGPSVFLFPPKPKDTLMISRTPEVTCVVVDVSHEDPEVKFNWYVDG<br>VEVHNAKTKPREEQYNSTYRVVSVLTVLHQDWLNGKEYKCKVSNKALPAPIE<br>KTISKAKGQPREPQVYTLPPSRDELTKNQVSLTCLVKGFYPSDIAVEWESNG<br>QPENNYKTTPPVLDSDGSFFLYSKLTVDKSRWQQGNVFSCSVMHEALHNHY<br>TQKSLSLSPGKTGGSGVDYPYDVPDYALD |
